# Genetic, biochemical, and molecular characterization of *Methanosarcina barkeri* mutants lacking three distinct classes of hydrogenase

**DOI:** 10.1101/334656

**Authors:** Thomas D. Mand, Gargi Kulkarni, William W. Metcalf

## Abstract

The methanogenic archaeon *Methanosarcina barkeri* encodes three distinct types of hydrogenase, whose functions vary depending on the growth substrate. These include the F420-dependent (Frh), methanophenazine-dependent (Vht), and ferredoxin-dependent (Ech) hydrogenases. To investigate their physiological roles, we characterized a series of mutants lacking each hydrogenase in various combinations. Mutants lacking Frh, Vht, or Ech in any combination failed to grow on H_2_/CO_2_, whereas only Vht and Ech were essential for growth on acetate. In contrast, a mutant lacking all three grew on methanol with a final growth yield similar to wild-type, produced methane and CO_2_ in the expected 3:1 ratio, but had a *ca.* 33% slower growth rate. Thus, hydrogenases play a significant, but non-essential, role during growth on this substrate. As previously observed, mutants lacking Ech fail to grow on methanol/H_2_ unless supplemented with biosynthetic precursors. Interestingly, this phenotype was abolished in the Δ*ech/*Δ*frh* and Δ*ech*/Δ*frh*/Δ*vht* mutants, consistent with the idea that hydrogenases inhibit methanol oxidation in the presence of H_2_, which prevents production of reducing equivalents needed for biosynthesis. Quantification of methane and CO_2_ produced from methanol by resting cell suspensions of various mutants supports this conclusion. Based on global transcriptional profiles, none of the hydrogenases are upregulated to compensate for loss of the others. However, transcript levels of the F420 dehydrogenase operon were significantly higher in all strains lacking *frh*, suggesting a mechanism to sense the redox state of F420. The roles of the hydrogenases in energy conservation during growth with each methanogenic pathway are discussed.

## IMPORTANCE

Methanogenic archaea are key players in the global carbon cycle due to their ability to facilitate the remineralization of organic substrates in many anaerobic environments. The consequences of biological methanogenesis are far reaching, with impacts on atmospheric methane and CO_2_ concentrations, agriculture, energy production, waste treatment and human health. The data presented here clarify the *in vivo* function of hydrogenases during methanogenesis, which in turn deepens our understanding of this unique form of metabolism. This knowledge is critical for a variety of important issues ranging from atmospheric composition to human health.

## INTRODUCTION

The ability to metabolize molecular hydrogen (H_2_) is a key metabolic feature in methanogenic Archaea (1). This trait is conferred by a class of enzymes known as hydrogenases, which catalyze the reversible oxidation of H_2_ coupled to various electron donors/acceptors (2, 3). At least five distinct types of hydrogenases are found in methanogenic archaea. These enzymes differ with respect to their redox partners, their cellular localization, and whether their activity is linked to production or consumption of membrane potential (4). Biochemical characterization of these diverse hydrogenases led to proposed functions for each enzyme class that differ substantially between methanogens with and without cytochromes (5).

Methanogens without cytochromes produce at least four types of hydrogenase; including (*i)* the electron-bifurcating Mvh hydrogenase, which couples oxidation of hydrogen to reduction of ferredoxin and a mixed coenzyme M-coenzyme B disulfide, (*ii)* the coenzyme F420-dependent hydrogenase, (*iii)* the [Fe] hydrogenase, which couples hydrogen oxidation to reduction of methenyltetrahydromethanopterin, and (*iv)* the ferredoxin-dependent, energy-converting hydrogenases (4). The first three are cytoplasmic enzymes, which supply the electrons needed to reduce CO_2_ to methane. The last is a membrane bound multi-subunit complex that couples hydrogenase activity to the production/consumption of the ion-motive force across the cell membrane (hence the designation as “energy-converting”). In non-cytochrome-containing methanogens, these energy-converting hydrogenases are believed to provide low-potential electrons, in the form of reduced ferredoxin, needed for anaplerotic reactions (6).

Methanogens with cytochromes, typified by *Methanosarcina barkeri*, encode a different set of hydrogenases that includes one cytoplasmic and two membrane-bound enzymes (Fig 1) (7). Like the non-cytochrome containing methanogens, *M. barkeri* produces a cytoplasmic, three-subunit F_420_-dependent hydrogenase known as Frh (for <u>F</u>420-<u>r</u>educing <u>h</u>ydrogenase). Frh is encoded by the four-gene *frhADGB* operon, which encodes the α, β and γ subunits (FrhA, FrhB and FrhG, respectively), along with the maturation protease FrhD (8). A second locus, *freAEGB*, encodes proteins that are 86-88% identical to FrhA, FrhB and FrhG, but lacks the gene for the maturation protease FrhD, instead encoding a small protein of unknown function (FrhE). It is not known whether the *fre* operon is capable of producing an active hydrogenase (8-11). A membrane-bound hydrogenase linked to the quinone-like electron carrier methanophenazine has, to date, been found only in *Methanosarcina* species. This enzyme, known as Vht because it was initially identified as a <u>v</u>iologen-reducing <u>h</u>ydrogenase, is encoded by the *vhtGACD* operon, which encodes the biochemically characterized enzyme comprised of VhtA and VhtG, along with a putative membrane-bound cytochrome, VhtC, that does not co-purify with the active enzyme, and a maturation protease, VhtD (7). As with the F420-reducing hydrogenase, a second locus that lacks the maturation protease is encoded in *M. barkeri* strains. This operon (*vhxGAC*) encodes proteins that display *ca.* 50% amino acid identity with those encoded by *vhtGACD.* Like *freAEGB*, it is not known whether the *vhx* operon produces an active hydrogenase (7). Finally, *M. barkeri* encodes a membrane-bound, ferredoxin-dependent <u>e</u>nergy-<u>c</u>onverting <u>h</u>ydrogenase (Ech) (12). This five-subunit enzyme complex is much simpler than, and only distantly related to, the energy-converting hydrogenases of the non-cytochrome methanogens, which typically contain more than a dozen subunits (3). Homologs of the electron bifurcating- and methenyltetrahydromethanopterin-reducing hydrogenases are not known to occur in cytochrome-containing methanogens.

**Fig 1.**
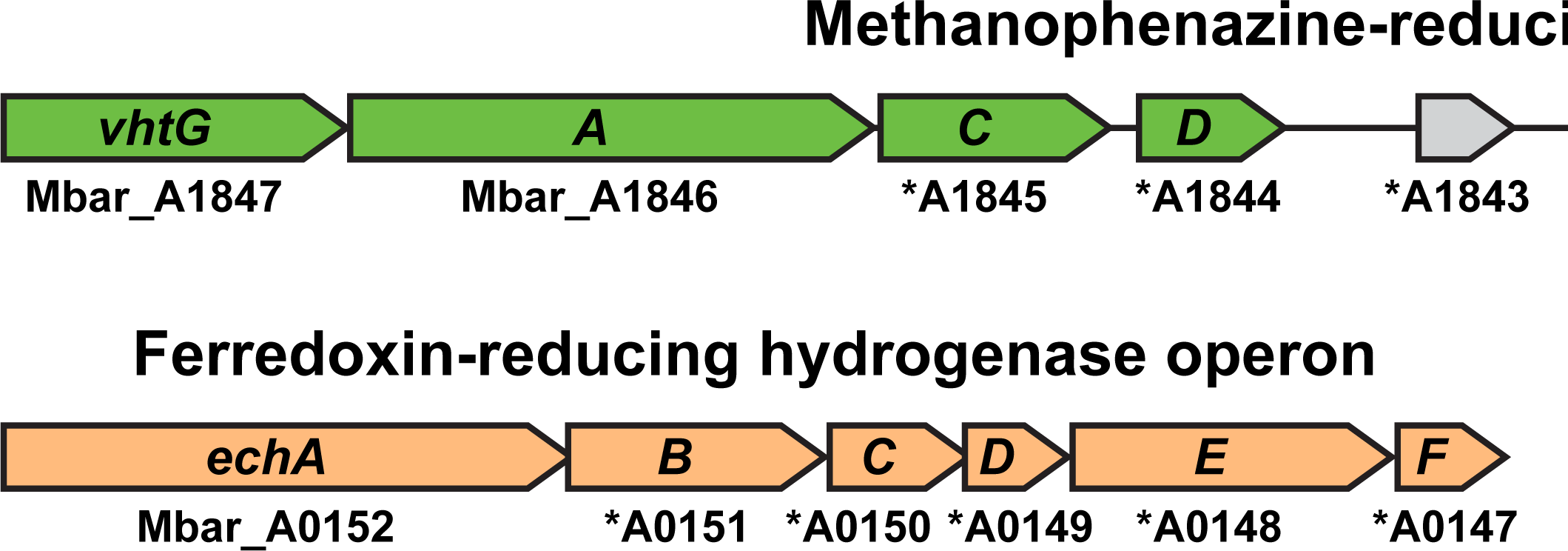
Hydrogenase operons of *Methanosarcina barkeri*. Three distinct types of hydrogenase are encoded by *M. barkeri*. Two potential methanophenazine-dependent hydrogenases are encoded by the adjacent *vhtGACD* and *vhxGAC* operons, while two potential F420-dependent hydrogenases are encoded by the unlinked *frhADGB* and *freAEGB* operons. The energy-converting, ferredoxin-dependent hydrogenase is encoded by the *echABCDEF* operon. Locus Tags are shown below each gene, in some cases the “Mbar_” prefix was omitted (shown by an asterisk) due to space constraints.

A key difference between the cytochrome-containing and non-cytochrome-containing methanogens is the ability of the former to use one-carbon (C-1) compounds and acetic acid, in addition to H_2_/CO_2_, as growth substrates. Catabolism of these chemically diverse substrates involves four distinct methanogenic pathways: the CO_2_ reduction pathway, the methyl reduction pathway, the methylotrophic pathway and the aceticlastic pathway (13, 14). While several of these pathways share common steps, they differ substantially with respect to involvement of key enzymes and the direction of metabolic flux during methane production (Fig 2). Surprisingly, it now appears that some *Methanosarcina* species (*e.g. M. barkeri*) use hydrogenases in each of the four pathways, regardless of whether external H_2_ is provided as a growth substrate (15).

**Fig 2.**
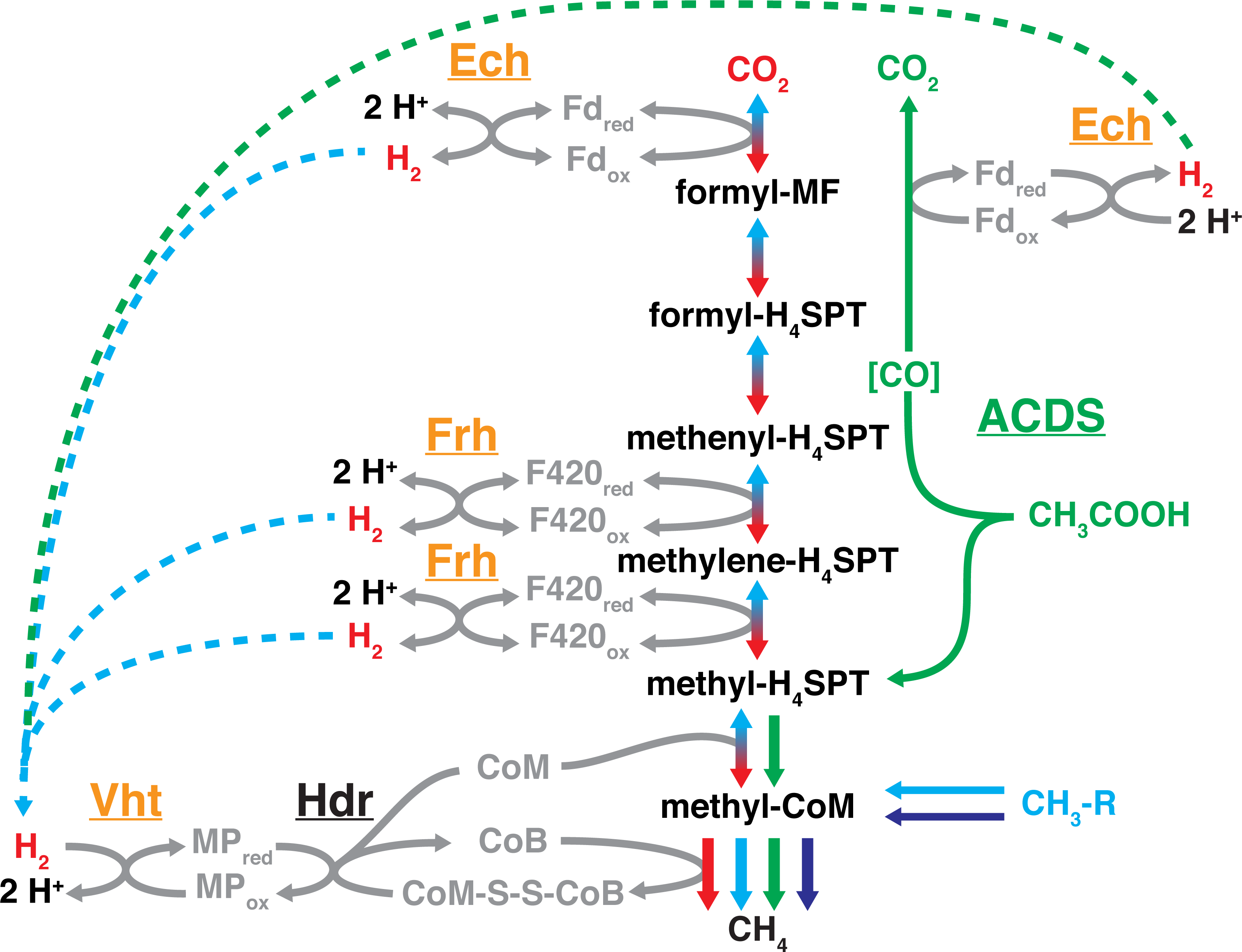
Four methanogenic pathways of *Methanosarcina*. Each pathway shares a common step in the reduction of methyl-CoM to methane; however, they differ in the route used to form methyl-CoM and in the source of electrons used for its reduction to methane. The CO_2_ reduction pathway (red arrows) involves reduction of CO_2_ to methane using electrons derived from the oxidation of H_2_, while the methylotrophic pathway (light blue arrows) involves disproportionation of C-1 substrates to methane and CO_2_. These two pathways share many steps, with overall metabolic flux in opposite directions (shown by red/light blue shaded arrows). In the aceticlastic pathway (green arrows), acetate is split into a methyl group and an enzyme-bound carbonyl moiety (shown in brackets) by the enzyme acetyl-CoA decarbonylase/synthase (ACDS). The latter is oxidized to CO_2_, producing reduced ferredoxin that provides electrons for reduction of methyl group to methane. Lastly, in the methyl reduction pathway (dark blue arrows) C-1 compounds are reduced to CH_4_ using electrons derived from H_2_ oxidation. Dashed lines depict diffusion of H_2_, which occurs during the transfer of electrons between oxidative and reductive portions of methylotrophic and aceticlastic pathways. The steps catalyzed by Frh, Vht, Ech hydrogenases are indicated. Note that in wild-type *M. barkeri* hydrogenases are involved in all four pathways. Abbreviations used are: Hdr, heterodisulfide reductase; MF, methanofuran; H_4_SPT, tetrahydrosarcinapterin; CoM, coenzyme M; CoB, coenzyme B; CoM-S-S-CoB, heterodisulfide of CoM and CoB; Fd_ox_/Fd_red_, oxidized and reduced ferredoxin; F420_ox_/F420_red_, oxidized and reduced cofactor F420; MP_ox_/MP_red_, oxidized and reduced methanophenazine.

During the CO_2_ reduction pathway, wherein CO_2_ is reduced to CH_4_ in a stepwise manner, hydrogenases produce the electron-donating cofactors required for four distinct reduction steps (Fig 2) (5). Initial reduction of CO_2_ to formyl-methanofuran requires reduced Fd (Fd_red_), which is produced by Ech. This reaction is dependent on ion-motive force because reduction of Fd by H_2_ is endergonic under physiological conditions (16). Subsequent reduction of methenyl-tetrahydrosarcinapterin (H_4_SPT) to methylene-H_4_SPT, and of methylene-H_4_SPT to methyl-H_4_SPT, requires reduced coenzyme F420 (F420_red_), which is supplied by Frh. Finally, reduction of a methyl group to methane using coenzyme B produces a heterodisulfide of coenzyme M and coenzyme B (CoM-S-S-CoB), which must be reduced to produce the free CoM and CoB needed for continued methanogenesis. This reaction is catalyzed by a membrane bound heterodisulfide reductase (HdrED), which uses the reduced form of a membrane-bound cofactor, methanophenazine, as the source of electrons (17). H_2_, in turn, serves as the reductant for generation of reduced methanophenzaine via membrane-bound Vht hydrogenase. Thus, all three types of hydrogenase are predicted to be required for growth via CO_2_ reduction, a conclusion that has been validated by the phenotypic analysis of single mutants (11, 15, 16).

In contrast, methanogenesis via the methyl reduction pathway is expected to require only Vht. In this pathway, methyl-CoM derived from C-1 compounds, such as methanol or methylamines, is directly reduced to methane using CoB as the electron donor. As with the CO_2_ reduction pathway, this produces a CoM-S-S-CoB disulfide that must be regenerated in a pathway requiring HdrDE and reduced methanophenazine, which is presumably generated by Vht (Fig 2). This idea is supported by the analysis of conditional *vht* mutants (15). Neither Frh nor Ech is required for methanogenesis in this model, a finding that is consistent with experimental data from Δ*frh* and Δ*ech* mutants (11, 16). Nevertheless, the *M. barkeri* Δ*ech* strain cannot grow via the methyl reduction pathway unless the media are supplemented with acetate and/or pyruvate. Thus, Ech plays an essential biosynthetic role under these conditions, which is probably the H_2_-dependent synthesis of reduced ferredoxin needed for synthesis of acetyl-CoA and pyruvate (16).

During aceticlastic methanogenesis, both the Ech and Vht hydrogenases play a critical role in methanogenesis. In this pathway, acetyl-CoA is split into methyl-H_4_SPT and enzyme-bound [CO] by the acetyl-CoA decarbonylase/synthase (ACDS) enzyme complex. CO is then further oxidized to CO_2_ with concomitant reduction of ferredoxin (12, 16). It is believed that the exergonic oxidation of ferredoxin by Ech produces H_2_ and contributes to the proton motive force by transferring protons across the membrane. The proton motive force is further enhanced by a putative H_2_ cycling mechanism, in which the H_2_ produced by Ech diffuses across the membrane, where it is oxidized by Vht to produce reduced methanophenazine. This unusual electron transport chain is completed when the reduced methanophenazine produced by Vht is used by HdrDE to regenerate free CoM and CoB from the CoM-S-S-CoB heterodisulfide (Fig 2) (18). Participation of these hydrogenases in the aceticlastic pathway is supported by mutagenic studies showing that *ech* and *vht* mutants do not grow with acetate as the sole substrate, regardless of whether biosynthetic precursors were supplied (15, 16).

Finally, all three types of hydrogenases are thought to be involved in methylotrophic methanogenesis via a H_2_ cycling mechanism similar to that described for aceticlastic growth (15). In this pathway, F420_red_ and Fd_red_, produced by the stepwise oxidation of methyl groups to CO_2_, are converted to molecular H_2_ by Frh and Ech, respectively (Fig 2). H_2_ then diffuses to the outer surface of the cell membrane where it is oxidized by Vht, releasing protons on the outside of the cell and contributing to the generation of an ion-motive force (Fig 3) (15). Nevertheless, *M. barkeri* Δ*frh* and Δ*ech* strains are capable of methylotrophic growth, indicating the presence of alternative pathways for transfer of electrons from F420_red_ and Fd_red_ to the electron transport chain (11, 16). The membrane-bound F420 dehydrogenase complex (Fpo) has been identified as the alternate mechanism of electron transfer from F420_red_ (11). This enzyme couples the exergonic reduction of methanophenazine by F420_red_, with the generation of proton motive force in a H_2_ independent manner. However, the *M. barkeri* Δ*frh* strain exhibits slower growth rates than wild type *M. barkeri*, showing that the H_2_-independent electron transport chain is less effective than electron transport via H_2_ cycling. The observation that Δ*fpo*/Δ*frh* double mutants are incapable of methylotrophic growth shows that additional electron transport routes are either not present, or not sufficient for methylotrophic growth (11). Mutants lacking Vht are inviable under all growth conditions, including methylotrophic growth, unless Frh is also removed. This phenotype is probably due the inability to recapture H_2_ produced in the cytoplasm, which causes redox imbalance and cell lysis (15).

**Fig 3.**
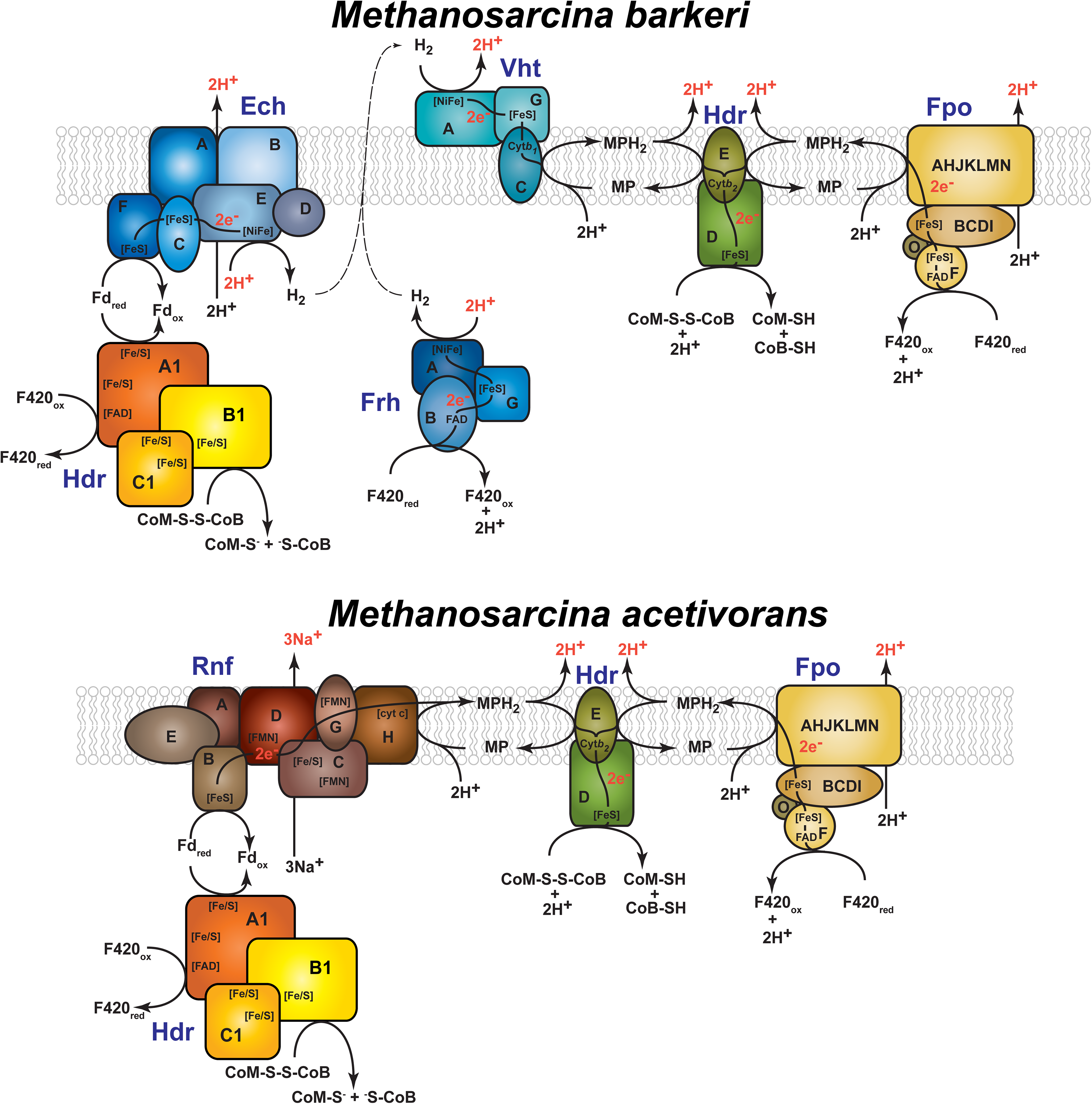
The branched electron transport systems of *Methanosarcina barkeri* and *Methanosarcina acetivorans.* During methylotrophic methanogenesis, *M. barkeri* can utilize H_2_-dependent or H_2_-independent electron transport systems. The H_2_-dependent pathway involves the use of hydrogenases Frh, Ech, and Vht in a H_2_ cycling mechanism to transfer electrons from F420_red_ or Fd_red_ to methanophenazine (MP). *M. acetivorans* does not utilize hydrogenases, and is therefore incapable of H_2_ cycling. Both organisms can conserve energy via a H_2_-independent pathway, wherein electrons are transferred from F420_red_ to MP by way of the F420 dehydrogenase (Fpo). Additionally, the electron transport system in *M. acetivorans* includes the Na^+^-translocating Rnf enzyme complex that serves as a Fd:MP oxidoreductase. Both membrane-bound electron transport systems converge at the reduction of the CoM-S-S-CoB heterodisulfide with electrons from reduced MP_red_ via the cytochrome-containing HdrDE enzyme. Protons (or Na^+^ in the case of Rnf) are translocated across the cell membrane in both systems, thereby conserving energy in the form of an ion motive force. An additional heterodisulfide reductase, HdrA1B1C1, has been proposed to function in the energy conservation pathways for both organisms. This electron bifurcating enzyme potentially reduces both the CoM-S-S-CoB heterodisulfide and F420 with electrons from Fd_red_ at a stoichiometric ratio of 2 Fd_red_ oxidized for the reduction of 1 CoM-S-S-CoB to CoM-SH/CoB-SH and reduction of 1 F420_ox_ to F420_red_. Abbreviations used are: Fd_ox_/Fd_red_, oxidized and reduced ferredoxin; MP/MPH_2_, oxidized and reduced methanophenazine; F420_ox_/F420_red_, oxidized and reduced F420; CoM, coenzyme M; CoB, coenzyme B; CoM-S-S-CoB, heterodisulfide of CoM and CoB; FeS, iron-sulfur cluster; FAD, flavin adenine dinucleotide; FMN, flavin mononucleotide; NiFe, nickel-iron active site of hydrogenases; cytb_1_/cytb_2_/cyt c, cytochromes.

Although the three *M. barkeri* hydrogenases have been studied *in vitro* and in certain mutants, a complete analysis of their role during growth on various substrates has yet to be reported. In this study, all five hydrogenase operons were systematically deleted in all viable combinations, and the physiological ramifications of these mutations examined by measuring growth, methanogenesis and hydrogenase activity on various growth substrates. We also performed global transcriptional profiling to assess the possibility that alternate electron transport chain components might be upregulated to compensate for the loss of specific hydrogenases. The data suggest that hydrogenases are not required for methylotrophic methanogenesis, but are essential for CO_2_ reductive, methyl reductive, and acetoclastic methanogenesis. Additionally, an inhibitory effect of H_2_ on the methyl oxidative pathway appears to be mediated by all three hydrogenases.

## RESULTS

### Construction of hydrogenase deletion mutants

To assess the role of the *M. barkeri* hydrogenases during growth on various substrates, we constructed mutants lacking the *frhADGB, freAEGB, vhtGACD, vhxGAC* and *echABCDEF* operons, individually and in all possible combinations (Fig S1). Because mutants lacking *vht* are only viable in Δ*frh* mutants (15), we also created a series of conditional mutants that have the *vht* promoter replaced by the synthetic P_*tet*_ promoter, which is expressed in the presence of tetracycline and tightly repressed in its absence (19). To simplify the isolation of strains lacking the adjacent *vht* and *vhx* loci, we constructed a mutant allele (denoted Δ*vht-vhx*) that deleted both operons along with two intervening genes that encode a putative peptidoglycan binding protein (Mbar_A1842) and an uncharacterized hypothetical protein (Mbar_A1843). The full set of strains containing deletions for hydrogenase operons in all possible combinations were successfully generated and verified by either Southern hybridization or PCR (Figures S2-S5).

### Characterization of growth phenotypes in hydrogenase deletion mutants

The generation time, growth yield and duration of lag phase for each mutant was determined by monitoring optical density during growth in a variety of media, providing clues as to the function of each hydrogenase during utilization of various carbon and energy sources (Table 1). With the exception of the Δ*fre* and Δ*vhx* mutations, which had no discernable phenotypes alone or in combination with other gene deletions, each of the mutations caused significant growth defects in one or more media.

**Table 1.**
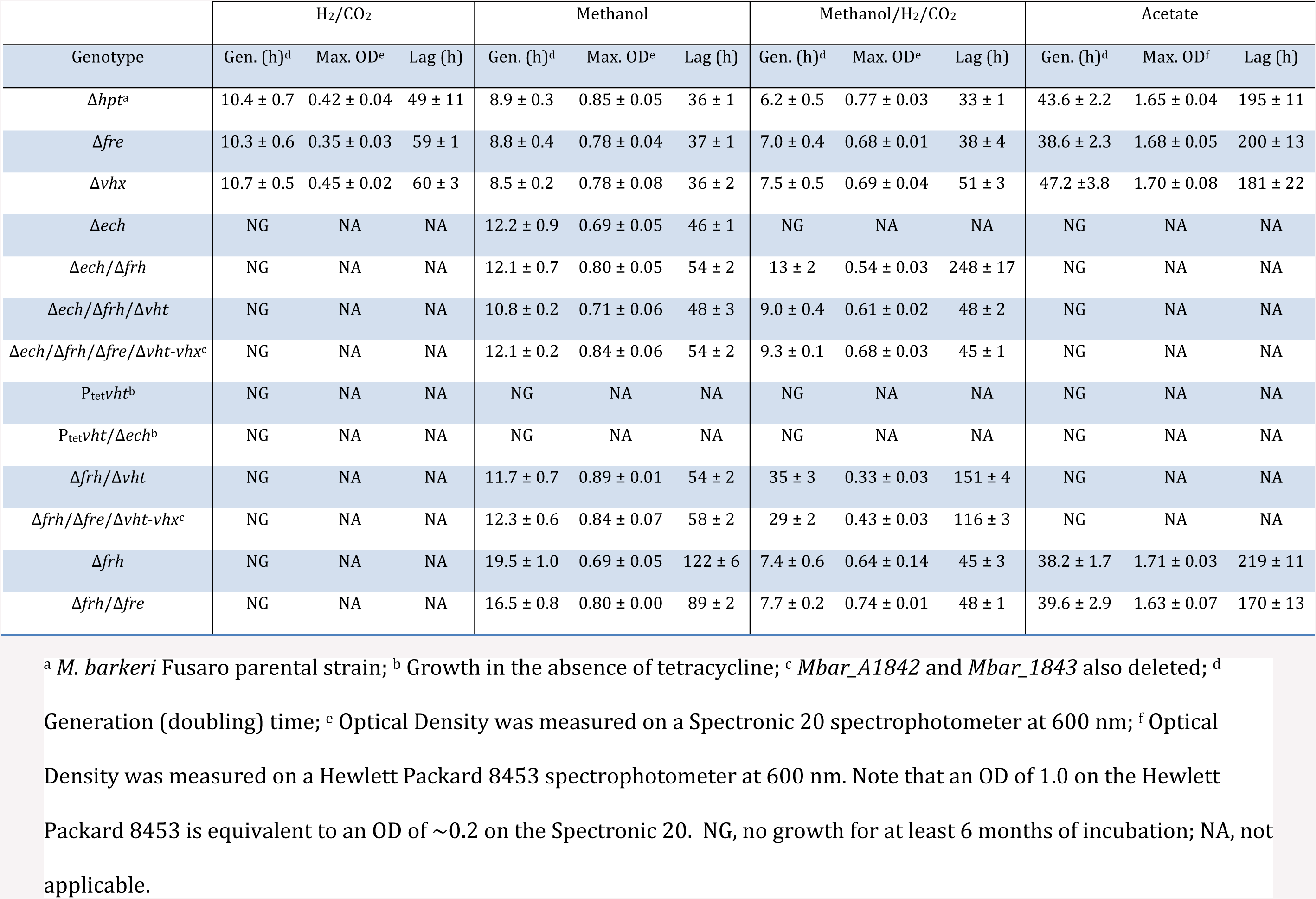
Growth of *M. barkeri* mutant strains on various methanogenic substrates.

Strains containing the Δ*ech* mutation were unable to grow in either H_2_/CO_2_ or acetate media, regardless of whether the other hydrogenase genes were deleted. However, with the exception of the P_*tet*_*vht/*Δ*ech* mutant discussed below, these mutants all grew on methanol medium, albeit with reduced growth rates. Consistent with previous reports (16), the Δ*ech* single mutant was unable to grow on unsupplemented methanol/H_2_/CO_2_ medium. Interestingly, the Δ*ech*/Δ*frh* double mutant regained the ability to grow in this medium, but with diminished rate and yield and the longest lag phase observed in any of our experiments. These phenotypes were substantially minimized in the Δ*ech*/Δ*frh*/Δ*vht* triple mutant, suggesting that both Frh and Vht inhibit methanol oxidation, which is needed to provide reducing equivalents for biosynthesis, when H_2_ is present.

As previously reported, we were unable to obtain a mutant lacking only *vht*, suggesting that loss of this locus is lethal in otherwise wild-type strains (15). This conclusion was supported by the phenotype of the P_tet_*vht* and P_*tet*_*vht/*Δ*ech* mutants, which were incapable of growth on any medium when tetracycline is absent (*i.e.* under repressing conditions). However, as previously noted, when cells were grown on methanol it was possible to delete the *vht* operon if *frh* was deleted first. The Δ*frh*/Δ*vht* strains, including ones that also carried an *ech* deletion, had methanol growth phenotypes similar to that of the wild type. Thus, hydrogenases are not required for growth on methanol, although *vht*-deficient strains are inviable in the presence of an active Frh hydrogenase. In contrast, strains lacking *vht* alone or in combination with other mutations were unable to grow on either H_2_/CO_2_ or acetate media. A more graded response was observed when various *vht* mutants were grown on methanol/H_2_/CO_2_. Accordingly, on this substrate combination, the *vht* single mutant was inviable, while the *Δfrh*/*Δvht* double mutant grew very poorly and the *Δech*/*Δfrh*/*Δvht* strain had phenotypes that were nearly wild-type. Again, these data are consistent with the idea that with certain mutant backgrounds Frh, Vht and Ech inhibit methanol oxidation in the presence of H_2_.

Finally, mutants lacking only the *frh* operon grew on three out of four substrates tested, failing to grow only on H_2_/CO_2_ medium. When methanol was the sole substrate, the Δ*frh* mutant had an extended lag phase and a generation time approximately double that of the parental strain. However, during growth on either acetate or methanol/H_2_/CO_2_ the growth phenotypes of this strain were equivalent to the parental strain, suggesting Frh enhances growth on methanol, but is not required for growth on the two latter substrate combinations.

### Methane and CO_2_ production by hydrogenase deletion mutants

To probe the underlying mechanisms behind the growth phenotypes, we also examined production of methane and CO_2_ by resting cell suspensions incubated with various substrates (Tables 2 & 3). The P_tet_*vht* and P_*tet*_*vht/*Δ*ech* mutants were not examined because they do not grow in any medium under non-inducing conditions. Similarly, we did not assay production of methane from acetate, because prior growth on acetate is required to induce the enzymes needed for this activity and most of the hydrogenase mutants are unable to grow under these conditions (20, 21).

**Table 2.** Production of CH_4_ and CO_2_ from resting cell suspensions of *M. barkeri* mutant strains.

| Genotype | H<sub>2</sub>/CO<sub>2</sub> |  | Methanol |  |  | Methanol/H<sub>2</sub> |  |  |
| --- | --- | --- | --- | --- | --- | --- | --- | --- |
|  | CH<sub>4</sub><sup>a</sup> | CO<sub>2</sub><sup>a</sup> | CH<sub>4</sub><sup>a</sup> | CO<sub>2</sub><sup>a</sup> | CH<sub>4</sub>: CO<sub>2</sub> | CH<sub>4</sub><sup>a</sup> | CO<sub>2</sub><sup>a</sup> | CH<sub>4</sub>: CO<sub>2</sub> |
| *Δhpt*<sup>b</sup> | 307 ± 24 | NA | 339 ± 6 | 118 ± 2 | 2.9:1 | 458 ± 6 | < 1 |  |
| *Δfre* | 316 ± 16 | NA | 339 ± 7 | 117 ± 2 | 2.9:1 | 442 ± 24 | < 1 |  |
| *Δvhx* | 328 ± 21 | NA | 329 ± 7 | 113 ± 3 | 2.9:1 | 453 ± 6 | < 1 |  |
| *Δech* | 5 ± 1 | NA | 331 ± 6 | 109 ± 2 | 3.0:1 | 460 ± 12 | < 1 |  |
| *Δech/Δfrh* | 3 ± 0 | NA | 328 ± 7 | 107 ± 2 | 3.1:1 | 440 ± 14 | 4 ± 1 | 110:1 |
| *Δech/Δfrh/Δvht* | 1 ± 0 | NA | 315 ± 5 | 93 ± 1 | 3.4:1 | 313 ± 5 | 92 ± 2 | 3.4:1 |
| *Δech/Δfrh/Δfre/Δvht-vhx*<sup>c</sup> | 1 ± 0 | NA | 314 ± 2 | 94 ± 1 | 3.3:1 | 315 ± 8 | 93 ± 3 | 3.4:1 |
| *Δfrh* | 34 ± 1 | NA | 310 ± 8 | 96 ± 2 | 3.2:1 | 442 ± 4 | < 1 |  |
| *Δfrh/Δfre* | 10 ± 2 | NA | 298 ± 13 | 98 ± 4 | 3.0:1 | 436 ± 8 | < 1 |  |
| *Δfrh/Δvht* | 6 ± 1 | NA | 311 ± 5 | 92 ± 1 | 3.4:1 | 225 ± 16 | 20 ± 1 | 11:1 |
| *Δfrh/Δfre/Δvht-vhx*<sup>c</sup> | 5 ± 0 | NA | 309 ± 6 | 92 ± 2 | 3.4:1 | 300 ± 8 | 19 ± 1 | 16:1 |

<sup>a</sup> μmol product observed; <sup>b</sup> *M. barkeri* Fusaro parental strain; <sup>c</sup> *Mbar\_A1842* and *Mbar\_1843* also deleted; NA, not applicable as produced CO<sub>2</sub> could not be differentiated from CO<sub>2</sub> that was added to the headspace. When relevant, ratio of CH<sub>4</sub>:CO<sub>2</sub> is shown. CH<sub>4</sub> and CO<sub>2</sub> quantities were below 4 ± 1 μmol for each strain when measured in cell suspensions without substrate.

**Table 3.** Rate of CH_4_ production^a^ from resting cell suspensions of *M. barkeri* mutant strains.

| Genotype | Methanol | Methanol & H<sub>2</sub> |
| --- | --- | --- |
| $\Delta hpt^b$ | 86 ± 7 | 132 ± 18 |
| $\Delta fre$ | 84 ± 1 | 123 ± 16 |
| $\Delta vhx$ | 71 ± 2 | 74 ± 12 |
| $\Delta ech$ | 57 ± 5 | 122 ± 7 |
| $\Delta ech/\Delta frh$ | 20 ± 1 | 27 ± 3 |
| $\Delta ech/\Delta frh/\Delta vht$ | 31 ± 2 | 30 ± 3 |
| $\Delta ech/\Delta frh/\Delta fre/\Delta vht-vhx^c$ | 19 ± 2 | 19 ± 2 |
| $\Delta frh$ | 23 ± 1 | 136 ± 9 |
| $\Delta frh/\Delta fre$ | 14 ± 1 | 87 ± 3 |
| $\Delta frh/\Delta vht$ | 38 ± 8 | 36 ± 4 |
| $\Delta frh/\Delta fre/\Delta vht-vhx^c$ | 32 ± 10 | 34 ± 13 |

<sup>a</sup> nmol methane produced min<sup>-1</sup> mg<sup>-1</sup>; <sup>b</sup> *M. barkeri* Fusaro parental strain; <sup>c</sup>
*Mbar\_A1842* and *Mbar\_1843* also deleted.

Consistent with their lack of growth phenotypes, the Δ*fre* and Δ*vhx* single mutations did not affect the levels of methane produced from any substrate tested. Neither did these mutations affect the ratio of methane/CO_2_ produced from methanol or from methanol/H_2_. However, the Δ*vhx* mutation slowed the rate of methane production. The Δ*fre* mutation also slowed the rate of methane production, but only when combined with the Δ*frh* mutation.

As seen in previous studies, Δ*ech* mutants produced only minor amounts of methane from H_2_/CO_2_ (<2% relative to the parental strain), but produced wild-type levels from both methanol and methanol/H_2_. During incubations with methanol, methane and CO_2_ were produced in a 3:1 ratio, consistent with disproportionation of the substrate via the methylotrophic pathway (Fig 2). Cell suspensions incubated with methanol and H_2_ produced only methane, showing that addition of hydrogen inhibits methanol oxidation. The rate of methane production by the Δ*ech* mutant was somewhat slower than wild-type using both methanol and methanol/H_2_. This rate was further reduced when the *ech* deletion was combined with mutations removing the *frh* or *vht* operons. Accordingly, the Δ*ech*/Δ*frh* double mutant produced methane nearly 5 times slower than the wild-type strain. Interestingly, this mutant produced a small amount of CO_2_ in addition to the wild-type level of methane, indicating a small amount of methanol was oxidized. When the *ech, frh,* and *vht* hydrogenases were deleted together, the quantity and stoichiometry of methane and CO_2_ production was identical to that observed on methanol alone.

The Δ*frh* single mutant produced similar levels and ratios of methane and CO_2_ relative to the parental strain with methanol or methanol/H_2_. When H_2_/CO_2_ was the substrate, methane production was reduced *ca.* 10-fold, but, significantly, not abolished. Combining the Δ*frh* mutation with deletions of *vht* and *ech* reduced methane production from H_2_/CO_2_ to negligible levels. In contrast, minimal affects on methane and CO_2_ production or stoichiometry were observed when methanol was the sole substrate. However, when combined with deletions of *vht* or *ech*, the Δ*frh* mutants produced significant levels of CO_2_ when incubated with methanol/H_2_, and the triple Δ*ech/*Δ*frh/*Δ*vht* mutant produced similar levels to that seen in assays incubated with methanol alone. Rates of methane production were substantially slower than wild-type for all Δ*frh* mutants.

### Enzyme activity in hydrogenase mutants

The hydrogenase activity for selected deletion mutants was measured in the forward direction (H_2_ oxidation) to allow estimation of the contributions of each enzyme to overall activity (Table 4).

**Table 4.** Hydrogenase activity of *M. barkeri* mutant strains.

| Genotype | Specific Activity<sup>a</sup> |
| --- | --- |
| $\Delta hpt^b$ | 11.10 ± 4.29 |
| $\Delta fre$ | 9.18 ± 4.38 |
| $\Delta vhx$ | 8.51 ± 3.93 |
| $\Delta ech/\Delta frh/\Delta vht$ | **0.01 ± 0.00<sup>d</sup>** |
| $\Delta ech/\Delta frh/\Delta fre/\Delta vht-vhx^c$ | **0.02 ± 0.00** |
| $\Delta ech$ | 17.13 ± 4.90 |
| $\Delta frh/\Delta vht$ | **0.42 ± 0.09** |
| $\Delta ech/\Delta frh$ | **2.60 ± 1.36** |
| $\Delta frh$ | **3.12 ± 1.89** |

<sup>a</sup>μmol H<sub>2</sub> oxidized min<sup>-1</sup> mg<sup>-1</sup>; <sup>b</sup> *M. barkeri* Fusaro parental strain; <sup>c</sup> *Mbar\_A1842* and
*Mbar\_A1843* also deleted; <sup>d</sup> values that are significantly different (p-value < 0.01)
than the $\Delta hpt$ parental strain are indicated in bold.

The hydrogenase activity of mutants lacking Fre or Vhx was not statistically different from the parental strain. Moreover, hydrogenase activity of the Δ*ech/*Δ*frh/*Δ*vht* mutant, which still encodes Fre and Vhx, was not statistically different than the Δ*ech/*Δ*frh/*Δ*fre/*Δ*vht-vhx* mutant that lacks all five hydrogenase operons. Thus, the *freAEGB* and *vhxGAC* operons do not, by themselves, produce detectable levels of hydrogenase. Because Fre and Vhx are essentially inactive, the hydrogenase levels in the Δ*frh/*Δ*vht* and Δ*ech/*Δ*frh* strains can be attributed solely to Ech and Vht, respectively. Accordingly, Ech has the lowest activity of the three hydrogenases, accounting for *ca.* 4% of total activity, and with Vht activity being *ca.* 6-fold higher. Consistent with this conclusion, deletion of *ech* did not significantly affect hydrogenase activity, whereas the Δ*frh* strain had drastically diminished activity as compared to the parental strain. Additionally, activity from the Δ*frh* strain, which encodes both Vht and Ech, is roughly equivalent to the combined activities of strains encoding only Vht or only Ech. Because strains expressing only Frh hydrogenase are inviable, the activity of this hydrogenase cannot be directly determined from a mutant strain. However, the relative contribution of Frh can be estimated from the hydrogenase activities of other mutants. Thus, by subtracting the activities of Vht and Ech from that of the parental strain, we estimate that roughly 75% of hydrogenase activity can be attributed to Frh.

### Effect of hydrogenase deletions on mRNA abundance

Our estimate of the relative activities of the individual hydrogenases assumes that the expression levels for each hydrogenase are unaffected by deletion of the others. To explicitly examine this possibility, we determined the global mRNA abundance profiles for each mutant using RNA seq (Table S4 and Dataset S1). Importantly, the RNA used in this analysis was isolated from the same cultures that were assayed for hydrogenase activity.

As expected, the mRNA levels for the deleted genes in each mutant were significantly and substantially lower than the parental strain, providing an important validation that the correct strains were used in these assays. Moreover, no significant differences in mRNA abundance from the parent were observed for the remaining hydrogenases in any strain, showing that the expression of individual hydrogenase operons is not regulated by the presence/absence of other hydrogenase genes. Thus, the hydrogenase activities found in the various mutants accurately reflect the combined activities of each enzyme in all strains.

Large numbers of *M. barkeri* genes showed significant changes in mRNA abundance in the hydrogenase mutants, relative to the parental strain. Accordingly, 2.7% of all genes were differently regulated in strains with one or two deleted hydrogenases, whereas the Δ*ech/*Δ*frh/*Δ*vht* and Δ*ech/*Δ*frh/*Δ*fre/*Δ*vht-vhx* strains showed in 17.4% and 22.5% differently regulated genes, respectively (Dataset S1). Of these, most encode proteins with unknown functions or with annotated functions that do not appear to be related to energy conservation. One exception was the F420 dehydrogenase (*fpo*), whose mRNA abundance increased significantly in all strains lacking *frh* (Table S4). This result suggests that the cell has a mechanism to sense the redox state of F420, which is altered upon deletion of *frh*.

## DISCUSSION

While fully consistent with the proposed functions of the *M. barkeri* hydrogenases, our phenotypic characterization of mutants lends new insight into the flexibility and interconnected nature of methanogenic metabolism. For example, reduction of CO_2_ to CH_4_ is expected to require three kinds of electron donors: Fd_red_, F420_red_, and reduced methanophenazine (1). Consistent with this idea, mutants lacking hydrogenases that reduce Fd (Ech), F420 (Frh) or methanophenazine (Vht) are unable to grow on H_2_/CO_2_. Thus, we were surprised to observe production of methane from H_2_/CO_2_ in cell suspensions of Δ*frh* mutants. Assuming this process involves the standard CO_2_ reduction pathway, this would require an alternative source of F420_red_ for reduction of methenyl- and methylene-tetrahydrosarcinapterin (Fig 2). Two alternative sources can be envisioned: first, Fpo could produce F420_red_ using reduced methanophenazine as the electron donor via reverse electron transport driven by proton motive force; second, a soluble heterodisulfide reductase could produce F420_red_ via electron bifurcation using CoM-S-S-CoB and Fd_red_ as substrates (as suggested in (22, 23)). In the former mechanism, reduced methanophenazine would be derived from H_2_ using Vht; in the latter, Fd_red_ would be derived from H_2_ via Ech. Interestingly, double mutants lacking Frh and either Ech or Vht produce much less methane than the Δ*frh* single mutant, thus both alternate pathways may contribute to this phenotype. The inability of the Δ*frh* mutant to grow on H_2_/CO_2_ suggests that this alternate methane-producing pathway does not provide sufficient energy for growth, or that it fails to provide an essential biosynthetic precursor.

Similar metabolic flexibility is seen during methylotrophic methanogenesis, which can occur via H_2_-dependent or -independent mechanisms (Figures 2, 3) (11, 12, 15, 16). We previously showed that a hydrogen cycling mechanism involving Frh and Vht is the preferred mode of electron transport in *M. barkeri*. Nevertheless, *M. barkeri* is also capable of methylotrophic growth in the absence of Frh and Vht (11, 15). Data reported here reveal that methylotrophic growth in *M. barkeri* is possible when all three hydrogenases are deleted. Thus, we have created an *M. barkeri* strain similar to *Methanosarcina acetivorans*, which has no detectable hydrogenase activity, but which grows well on methylotrophic substrates (7, 14). During methylotrophic growth in *M. acetivorans,* an electron transport chain comprised of Fpo and HdrDE is used to capture energy from F420_red_ produced by the oxidative branch of the methanogenic pathway (Figures 2, 3). Our genetic analyses suggest that Fpo is also used to metabolize F420_red_ in *M. barkeri* Δ*frh/*Δ*vht* mutants (11, 15). The oxidative branch of the methylotrophic pathway also produces Fd_red_, which in *M. acetivorans* is oxidized by membrane-bound, ion-pumping Fd_red_:methanophenazine oxidoreductase known as Rnf (24). However, because *M. barkeri* does not encode Rnf, this energy-conserving electron transport pathway in not available to the Ech mutants characterized here. Thus, an alternative Fd_red_:heterodisulfide oxidoreductase system must exist to allow growth of these mutants on methanol. It has been suggested that this alternate Fd_red_:heterodisulfide oxidoreductase activity is catalyzed by a cytoplasmic, electron-bifurcating heterodisulfide reductase (HdrABC), similar to the electron-bifurcating heterodisulfide reductase of non-cytochrome containing methanogens (Fig 3) (22). Biochemical data from a homologous *M. acetivorans* enzyme supports this possibility (23).

Interestingly, the stoichiometry of methane and CO_2_ produced from methanol/H_2_ in Δ*frh/*Δ*vht* and Δ*frh/*Δ*fre/*Δ*vht-vhx* mutants lends additional support for an alternate Fd_red_:heterodisulfide oxidoreductase. These strains, which encode only Ech, might be expected to disproportionate methanol to CH_4_ and CO_2_ in a 3:1 ratio, as was seen in the strains lacking all three active hydrogenases. Instead, they produced CH_4_ and CO_2_ at an approximate ratio of 10:1, suggesting that a substantial portion of methanol was reduced directly to CH_4_ using electrons obtained by H_2_ oxidation. Because Ech is the sole remaining hydrogenase in these strains, electrons from Fd_red_ must be involved in this process.

H_2_ also inhibits oxidation of methanol when both substrates are present, via a mechanism that is clearly mediated by hydrogenase activity. Accordingly, in the presence of H_2_, methanol is solely reduced to methane by cell suspensions of strains that contain all three hydrogenases, while it is disproportionated to methane and CO_2_ in a 3:1 ratio when all three are absent. The hydrogenase-mediated inhibition of methanol oxidation is graded, with Vht having the largest effect and Ech the least.Similarly, Ech mutants are only able to grow on methanol/H_2_ when supplemented with acetate and pyruvate, which has been interpreted to mean they cannot produce the reducing equivalents needed for biosynthesis by oxidizing methanol to CO_2_ (16). We showed here that this effect is alleviated by deletion of genes encoding Frh and Vht, with the Δ*vht* mutation having a much larger effect. Because protein synthesis was blocked by addition of puromycin in the cell suspension experiments, these effects cannot have been mediated by changes in the concentration of enzymes in the methanogenic pathways. Moreover, because inhibition requires the hydrogenase enzymes to be present, it is likely that a product of the enzymatic reaction mediates inhibition: namely the reduced enzyme cofactors. Thus, in the presence of high H_2_ partial pressures and the appropriate hydrogenase, we would expect the levels of oxidized methanophenazine, F420_ox_ and Fd_ox_ to be kept at very low levels. Interestingly, the graded inhibition in response to loss of Vht, Frh and Ech mimics the thermodynamics of the hydrogenase reactions, with methanophenazine being the most energetically favorable electron acceptor and Fd being the least. This suggests at least two possible mechanisms that might account for the inhibition: *i)* allosteric inhibition or covalent inactivation of a key enzymatic step in the oxidative branch of the pathway could be triggered by one or more reduced cofactors, or *ii)* simple changes in the availability of F420_ox_ and Fd_ox_, which are needed for three discrete steps in the oxidative branch of the methyloptrophic pathway (Fig 2). Note that in the second mechanism, the major inhibitory effect of Vht on methyl oxidation can only be explained if high levels of reduced methanophenazine influence the levels of F420_ox_, which could occur by changing the equilibrium of the Fpo reaction (Fig 2, 3).

In addition to affecting flux through methanogenic pathways, the levels of reduced or oxidized cofactors may be used as a sensory input to modulate gene regulation. Transcriptional profiling of hydrogenase mutants showed that in all strains lacking *frh*, the *fpo* operon was significantly up-regulated. Without Frh, Fpo is solely responsible for the F420_red_:methanophenazine oxidoreductase activity required to transfer electrons from the oxidative to reductive portions of the methylotrophic electron transport pathway. Elevated abundance of *fpo* mRNA in Δ*frh* strains indicates that the cell has a mechanism to sense and respond to F420 redox imbalance. A previous study identified MreA as a global regulator in *Methanosarcina* with the ability to bind and repress the *fpo* promoter region during aceticlastic growth (25). This regulator was shown to affect gene expression based on growth substrate, however the mechanism and sensory input are unknown. Systems for gene regulation based on detected redox imbalance of F420 and other electron carriers are a potential source for future studies.

The levels of hydrogenase activity for the three enzyme types have significant ramifications for the hydrogen cycling model on energy conservation (15). We have shown that Δ*vht* mutations are lethal when Frh is present, but not when it is absent. Moreover, when *vht* expression is turned off using a regulated promoter, cell lysis is concomitant with H_2_ accumulation, implying that the inability to recapture H_2_ produced in the cytoplasm is responsible for the lethal phenotype. With this in mind, it seems clear that the cytoplasmic activities of Frh must be carefully balanced against the periplasmic activity of Vht. Interestingly, our data show that Frh activity is ca. 3-fold higher than that of Vht. Thus, it appears that ability of Frh to produce H_2_ is much higher than the ability of Vht to take it up. We recognize that our assays were not conducted with the native substrates (which are not commercially available), therefore we approximated the *in vivo* activity of each enzyme based on available literature values in which a variety of natural and artificial cofactors were used (Table S5). These data suggest that the relative activities of Frh and Vht are more similar than our assay data suggest, with Frh activity ca. 1.5-fold higher than Vht. While this extrapolation must be interpreted with caution, it still suggests that Frh capacity is higher than that of Vht. In this regard, both Vht and Ech are coupled to ion-motive force; thus, activity in whole, metabolically active cells could be substantially different.

Finally, unlike Frh and Vht, Fre and Vhx are not able to provide sufficient levels of F420_red_ and reduced methanophenazine, respectively, for growth via CO_2_ reduction. Additionally, the Δ*ech/*Δ*frh/*Δ*vht* strain that only encoded for the Fre and Vhx hydrogenases had no detectable hydrogenase activity. This could be due to low expression of *fre* and *vhx* operons, absence of post-translational processing, mutations in structural or catalytic residues or some combination of these (7). Analysis of RNA sequencing data from wild type *M. barkeri* grown methylotrophically indicated the abundance mRNA for *fre* approximately 50-lower than *frh* (Dataset S1), similar to the relative abundance observed by Vaupel and Thauer (8). Additionally, the abundance of *vhx* transcripts was more than 200-fold lower than those of *vht*. We note that our enzymatic assays would have easily detected hydrogenase activity at levels 200-fold lower than we observed for the strains encoding only *vht*. Thus, poor gene expression cannot explain the lack of activity in strains expressing only Fre of Vhx. Hydrogenases require several maturation steps to become active enzymes, including processing by the maturation proteases encoded by the *frhD* and *vhtD* genes. Thus, it remains possible that Fre and Vhx could encode active enzymes if FrhD and VhtD are *trans*-acting maturation proteases. Given that the mutants characterized here removed the entire *frh* and *vht* operons, our data do not address this possibility.

## MATERIALS AND METHODS

### Media and growth conditions

*Methanosarcina* strains were grown as single cells (26) at 37 °C in high salt (HS) broth medium (27) or on agar-solidified medium as described (28). Growth substrates provided were methanol (125 mM in broth medium and 50 mM in agar-solidified medium) or sodium acetate (120 mM) under a headspace of N_2_:CO_2_ (80:20 v/v) at 50 kPa over ambient pressure, H_2_:CO_2_ (80:20 v/v) at 300 kPa over ambient pressure, or a combination of methanol plus hydrogen. Cultures were supplemented as indicated with 0.1% yeast extract (YE), 0.1% casamino acids (CAA), 10 mM sodium acetate, 10 mM pyruvate or 100 mM pyruvate. Puromycin (CalBioChem, San Diego, CA) was added at 2 mg ml^-1^ for selection of the puromycin transacetylase (*pac*) gene (29). 8-Aza-2,6-diaminopurine (8-ADP) (Sigma, St Louis, MO) was added at 20 mg ml^-1^ for selection against the presence of *hpt* (29). Tetracycline (Tc) was added at 100 mg ml^-1^ to induce the tetracycline-regulated P*mcrB*(*tet*O1) promoter (19). Standard conditions were used for growth of *Escherichia coli* strains (30) DH5α/λ-*pir* (31) and DH10B (Stratagene, La Jolla, CA), which were used as hosts for plasmid constructions.

### DNA methods and plasmid constructions

Standard methods were used for plasmid DNA isolation and manipulation in *E. coli* (32). Liposome mediated transformation was used for *Methanosarcina* as described (33). Genomic DNA isolation and DNA hybridization were as described (27, 28, 34). DNA sequences were determined from double-stranded templates by the W.M. Keck Center for comparative and functional genomics, University of Illinois. Plasmid constructions are described in Tables S1 and S2.

### Strain construction in *M. barkeri*

The construction and genotype of all *Methanosarcina* strains is presented in Table S3. Hydrogenase encoding genes were deleted sequentially in a specific order (Figure S1) because certain hydrogenase deletion mutants are only viable when other hydrogenase genes are deleted first (15). To simplify isolation of strains that lack hydrogenase operons *vhxGAC* and *vhtGACD*, the genes between the two operons (*Mbar_A1842* and *Mbar_A1843*) were also deleted (Figure 1). All mutants were confirmed by either PCR or DNA hybridization (Figures S2-S5).

### Determination of growth characteristics

For growth rate determinations, cultures were grown on methanol or methanol plus H_2_/CO_2_ (Δ*frh* and Δ*frh/*Δ*fre*) to mid-log phase (optical density at 600 nm [OD_600_] *ca.* 0.5). Approximately 3% inoculum of the culture (or 10%, in case of acetate) was then transferred to fresh medium in at least four replicates and incubated at 37 °C. Growth was quantified by measuring OD_600_. With the exception of samples grown on acetate, all OD_600_ were measured with a Spectronic 20 spectrophotometer (Thermo Fisher Scientific, Waltham, MA); those grown on acetate were measured with a Hewlett Packard 8453 spectrophotometer (Agilent,Santa Clara, CA). Note that an OD of 1.0 on the Hewlett Packard 8453 is equivalent to an OD of ˜ 0.2 on the Spectronic 20. Generation times were calculated during exponential growth phase and lag phase was defined as the time required to reach half-maximal OD_600_.

### Cell suspension experiments

Cells grown on methanol or methanol plus H_2_/CO_2_ (Δ*frh* and Δ*frh/*Δ*fre*) were collected in late exponential phase (OD_600_ = 0.6-0.7) by centrifugation at 5,000 x g for 15 minutes at 4 °C. The cells were washed once with anaerobic HS PIPES buffer (50 mM PIPES at pH 6.8, 400 mM NaCl, 13 mM KCl, 54 mM MgCl_2_, 2 mM CaCl_2_, 2.8 mM cysteine, 0.4 mM Na_2_S) and resuspended in the same buffer to a final concentration of 10^9^ cells/ml. Cells were counted visually using the Petroff-Hausser counting chamber (Hausser Scientific, PA). All assay mixtures contained 2 ml of the suspension and were conducted under strictly anaerobic conditions in 25 ml Balch tubes sealed with butyl rubber stoppers using 250 mM methanol as the methanogenic substrate under a headspace of N_2_, H_2_, or H_2_/CO_2_ (80/20%) at 250 kPa over the ambient pressure, as indicated. Puromycin (20 μg/ml) was added to prevent protein synthesis. Cells were held on ice until initiation of assay by transfer to 37 °C. For rate determination, gas phase samples were withdrawn at various time points and assayed for CH_4_ by gas chromatography at 225 °C in a Hewlett Packard gas chromatograph (5890 Series II) equipped with a flame ionization detector. The column used was stainless steel filled with 80/120 Carbopack^TM^ B/3% SP^TM^-1500 (Supelco, Bellefonte, PA) with helium as the carrier gas. For total CH_4_ and CO_2_ production, assays were incubated at 37 °C for 36 hours prior to withdrawal of gas phase samples for analysis by GC at 225 °C in a Hewlett Packard gas chromatograph (5890 Series II) equipped with a thermal conductivity detector. A stainless steel 60/80 Carboxen-1000 column (Supelco, Bellefonte, PA) with helium as the carrier gas was used. Total cell protein was determined using the Bradford method (35) after 1 ml of the cells was lysed by resuspension in ddH_2_0 with 1 mg/ml RNase and DNase.

### Hydrogenase assays

Strains were grown at 37 °C in HS medium supplemented with 125 mM methanol and cells were harvested from 10 ml mid-exponential phase culture at 1228 x *g* for 15 min in an IEC MediSpin (Needham Heights, MA) benchtop centrifuge. Preparation of cell extract was performed in an anaerobic chamber under an atmosphere of H_2_/N_2_ (4/96%). Cells were washed once in 10 ml HS-MOPS [2 mM dithiothreitol (DTT), 400 mM NaCl, 13 mM KCl, 54 mM MgCl_2_, 2 mM CaCl_2_, 50 mM MOPS, pH 7.0] and lysed in 1 ml lysis buffer (2 mM DTT, 0.5% *n*-dodecyl β-D-maltoside, *ca.* 50 Kunitz units bovine pancreas deoxyribonuclease I, 50 mM MOPS, pH 7.0) on ice for 30 min. Enzyme-containing supernatant was separated from cell debris by centrifugation at 13600 x *g* for 2.5 min (Fisher Scientific Micro Centrifuge Model 235C, Waltham, MA). Protein concentration was measured via the Bradford method (35).

Assays were performed anaerobically in 1.7 ml quartz cuvettes sealed with rubber stoppers. A total reaction volume of 1 ml was used, and included cell extract mixed with 50 mM MOPS buffer (pH 7.0) containing 2 mM DTT and 2 mM benzyl viologen (BV). The cuvette headspace was pressurized to 30 kPa with 100% H_2_ after being flushed for 2 min. Cuvettes with reaction mixture were pre-warmed to 30 °C before the reaction was initiated by the addition of BV. Hydrogenase activity was determined by quantifying the change in absorbance of BV at 578 nm (extinction coefficient, 8.65 cm^-1^ mM^-1^) with a Cary 50 UV-Vis Spectrophotometer (Agilent, Santa Clara, CA). One unit (U) of hydrogenase activity was defined as the oxidation of 1 μmol H_2_ per minute, based on the fact that 2 μmol BV are reduced for each H_2_ oxidized. A minimum of three independent measurements from biological replicates was performed for each strain.

### RNA sequencing

Immediately prior to cell harvest for hydrogenase assays, 2.5 ml of the same culture was harvested for total RNA isolation. An equal volume of TRIzol reagent (Ambion, Carlsbad, CA) was added to the culture to lyse cells and samples were incubated at room temperature for 5 min. RNA was then isolated with a Direct-zol RNA MiniPrep kit from Zymo Research (Irvine, CA) according to the manufacturer’s directions. RNA samples were stored at −80 °C.

To increase coverage of mRNA during sequencing, rRNA was removed from samples via subtractive hybridization. The method of Stewart et al. (36) was utilized with the following modifications. Templates for 16S and 23S rRNA probes were generated by PCR from strain WWM85 with primers 16SFor, T716SRev, 23SFor, and T723SRev. *In vitro* transcription with the MEGAscript High Yield Transcription kit (Ambion) was used for the production of biotinylated antisense rRNA probes from 400 ng of the purified PCR products in separate reactions. After removal of template with DNAseI, probes were purified with the Zymo Research RNA Clean & Concentrator kit. Hybridization reactions (30 μl) for each sample contained the following: 20% formamide, 1X SSC buffer, 20 U SUPERase inhibitor, 2 μg total RNA, 4 μg 16S probe, and 4 μg 23S probe. Reactions were denatured at 70 °C for 10 min, ramped down to 25 °C (−0.1 °C sec^-1^), and incubated at room temperature for 10 min. rRNA hybridized to biotinylated probe was removed via streptavidin-coated magnetic beads (New England Biolabs, Ipswich, MA). Beads (500 μl per sample) were washed twice with 500 μl 1X SSC buffer prior to the addition of hybridized RNA sample diluted to 250 μl in 1X SSC buffer with 20% formamide. Samples were incubated for 1 hour at room temperature with gentle shaking before separation of beads on a magnetic rack. The supernatant was removed, beads were washed with 250 μl 1X SSC, and supernatant and wash were pooled and cleaned with the Zymo Research RNA Clean & Concentrator kit.

Preparation and sequencing of RNAseq libraries was performed at the Roy J. Carver Biotechnology Center at the University of Illinois at Urbana-Champaign.Libraries were made with the TruSeq Stranded mRNA Sample Prep kit, sequenced with a HiSeq 2000 using the TruSeq SBS v3 kit, and processed with Casava 1.8.2, all per the manufacturer’s directions (Illumina, San Diego, CA). All sequencing data was further processed and analyzed as previously described (37) with CLC Genomics Workbench 7 (Qiagen). Within this program, the Empirical analysis of Differential Gene Expression (EDGE) tool was used for statistical analysis (38). Differently regulated genes were considered significant when up- or down-regulated at least 3-fold with a p-value ≤ 0.05. Three biological replicates were sequenced and analyzed for each strain. Raw and processed data have been deposited in the Gene Expression Omnibus (GEO) under the accession number GSE98441.

## ACKNOWLEDGEMENTS

The authors acknowledge the Division of Chemical Sciences, Geosciences, and Biosciences, Office of Basic Energy Sciences of the U.S. Department of Energy through Grant DE-FG02-02ER15296 for funding of this work.

